# CRISPR-typing PCR (ctPCR), a new Cas9-based DNA detection method

**DOI:** 10.1101/236588

**Authors:** Qiao Wang, Beibei Zhang, Xinhui Xu, Feifei Long, Jinke Wang

**Affiliations:** State Key Laboratory of Bioelectronics, Southeast University, Nanjing 210096, China; Nanjing Foreign Language School, Nanjing 210096, China

**Author notes:** These authors contributed equally to this study (Qiao Wang, Beibei Zhang). Correspondence should be addressed to Jinke Wang.

**Keywords:** CRISPR, PCR, nucleic acid, detection

## Abstract

This study develops a new method for detecting and typing target DNA based on Cas9 nuclease, which was named as ctPCR, representing Cas9/sgRNA- or CRISPR-typing PCR. The technique can detect and discriminate target DNA easily, rapidly, specifically, and sensitively. This technique detects target DNA in three steps: (1) amplifying target DNA with PCR by using a pair of universal primers (PCR1); (2) treating PCR1 products with a process referred to as CAT, representing Cas9 cutting, A tailing and T adaptor ligation; (3) amplifying the CAT-treated DNA with PCR by using a pair of general-specific primers (gs-primers) (PCR2). The technique was verified by detecting HPV16 and HPV18 L1 gene in 13 different high-risk human papillomavirus (HPV) subtypes. The technique was also detected two high-risk HPVs (HPV16 and HPV18) in cervical carcinoma cells (HeLa and SiHa) by detecting the L1 and E6/E7 genes, respectively. In this method, PCR1 was performed to determine if the detected DNA sample contained the target DNA (such as virus infection), while PCR2 was performed to discriminate which genotypic target DNA was present in the detected DNA sample (such as virus subtypes). With these proof-of-concept experiments, this study provides a new CRISPR-based DNA detection and typing method.

## Introduction

DNA detection and genotyping is always of importance to the basic researches and various detecting and diagnostic applications. Therefore, the DNA detection and genotyping techniques have been attracting increasing attention, which promotes the advance of technique development. Summarily, there are mainly three classes of widely used DNA detection and genotyping techniques. The first is various techniques based on polymerase chain reaction (PCR). PCR has been the most popular DNA detection and genotyping technique. PCR-based DNA detection and genotyping mainly relies on the design of highly specific PCR primers and multiple PCR amplification. PCR detection could be realized by the traditional PCR (tPCR), the quantitative PCR (qPCR), and the recently developed digital PCR^1^. Q-PCR has been highly popularized in almost all research, detection and diagnostic labs due to its obvious advantages such as real time and high sensitivity. Now, the more accurate digital PCR has already been developed, which has its great potential and advantages to be used as clinical detection tool ^2,3^. However, the PCR techniques are limited by the difficulty of multiple amplification and designing highly specific primers that can discriminate highly-related genotypes. In addition to PCR techniques, various DNA hybridization techniques such as DNA microarray has also been widely used to detect and type DNA ^4,5^. However, due to its costly equipment, complicate detection process and unavoidable nonspecific hybridization, DNA microarray technique is unable to become a routine DNA detection and genotyping tool as PCR. DNA sequencing is another efficient DNA detection and genotyping technique ^6,7^. Especially, with the advent of next-generation sequencing (NGS) techniques ^8,9^, the NGS platforms such as Illumina NovaSeq become increasingly available DNA sequencing tools. Nevertheless, due to their current costly equipment and chemical reagents, they are still unavailable to the routine research, detection and diagnosis as PCR. Therefore, in comparison, PCR is still a most convenient, cost-effective, and ready platform for DNA detection and genotyping, especially if its primer designing limitation is overcome.

The clustered regularly interspaced short palindromic repeat (CRISPR) were firstly found in the genome of *Escherichia coli* (*E.coli*) by Ishino et al in 1987 ^10^ and defined by Jansen et al in 2002 ^n^. Now, it is known that CRISPR system includes three different types (Type I, II and III) ^12,13^. Type I and III systems consist of multiple Cas proteins ^14^, while Type II system requires only one Cas protein, Cas9 ^15^. Cas9 can be associated with CRISPR-related RNA (crRNA) and trans-activating crRNA (tracrRNA) ^15^. TracrRNA is able to activate Cas9 nuclease and crRNA is complementary to 20 nucleotide sequence of target DNA. The latter thus determines the specificity of CRISPR-Cas9 system. The crRNA-guided Cas9 nuclease can bind to the target DNA neighbored by a protospacer adjacent motif (PAM) and cut two strands of target DNA at the same position of three bases upstream of PAM sequence (NGG) ^15,16^. The tracrRNA and crRNA were subsequently integrated into one RNA named as single-guided RNA (sgRNA) ^17^, which greatly simplified the application of type II CRISPR system. Cas9 is directed to cleave the target DNA by sgRNA ^18^. Currently, due to its simplicity and efficiency, CRISPR-Cas9 system is now widely being used to edit gene by countless researchers ^19^. In addition, the dead Cas9 (dCas9), which is inactivated to lose nuclease activity but fused with a gene transcriptional activation domain (AD) or inhibition domain (ID), has been widely used to regulate endogenous gene expression as a new kind of artificial transcription factor ^20^.

Despite Cas9/sgRNA has been widely applied to gene editing and regulation, it has not been fully exploited to develop nucleic acids detection techniques. Nevertheless, due to the sequence-specific DNA cutting capability (even single base distinction ^21^), Cas9/sgRNA has great potential to detect and type DNA. Recently, CRISPR-Cas9 system had been used to type American and African Zika viruses in a RNA in vitro transcription-based Zika virus detection technique ^22^. Given the high specificity of CRISPR-based tools, CRISPR-Cas9 can discriminate viral strains at single-base resolution ^22^, which is beneficial for detecting and typing the orthologous and homologous bacteria and viruses even at single nucleotide polymorphism level. Other CRISPR systems, such as Cas13a/C2c2 of type III, have been recently used to develop Zika virus detection technique that has ultrahigh sensitivity (viral particles down to 2 aM) ^23^. These studies indicate that CRISPR systems have great potential and advantages to be exploited to develop nucleic acid detection techniques. However, in the currently reported Cas9-based nucleic detection method ^22^, Cas9/sgRNA system was used to type RNAs by cutting the DNAs that were firstly reversely transcribed from the detected RNAs and then transferred into double-stranded DNA by a second strand synthesis. The Cas9/sgRNA system therefore has not been exploited to directly detect and type various DNA targets, which is the main object of routine nucleic acid detection.

In this study, we developed a new method for detecting target DNAs based on Cas9 nuclease, which was named as ctPCR, representing the Cas9/sgRNA- or CRISPR-typing PCR. In ctPCR detection, the target DNA was firstly amplified by a first-round PCR (PCR1) by using a pair of universal primers. The PCR1 products were then successively digested by Cas9 in complex with a pair of sgRNAs, tailed by an adenine (A), and ligated by a T adaptor. Finally, the treated PCR1 products were amplified by a second-round PCR (PCR2) using a pair of general-specific primers (gs-primers). PCR1 was used to find if the DNA sample contained the target DNA (such as virus infection), while PCR2 was used to discriminate which genotype of DNA presents in the DNA sample (such as virus subtype). This study demonstrated that ctPCR could detect the L1 genes of HPV16 and HPV18 from 13 different high-risk human papillomavirus (HPV) subtypes. This study also indicated that ctPCR could detect two high-risk HPVs (HPV16 and HPV18) in the human cervical carcinoma cells (HeLa and SiHa) by detecting both L1 and E6/E7 genes. By performing these proof-of-principle detections, this study developed a new CRISPR-based PCR method for detecting and typing DNA. This method takes advantages of PCR but eliminates hard primer designing by combining CRISPR with PCR. This method realized rapid DNA detection and typing without depending on hybridization and sequencing.

## Experimental section

### Construction of sgRNA expression plasmid

A pair of primers was firstly synthesized to amplify the prokaryotic Cas9 gene sequence with PCR from pCas9 (Addgene), in which the forward primer contained the J23100 promoter plus RBS sequences (total 88 bp). Then the prokaryotic Cas9 gene sequence was amplified from pCas9 (Addgene) by using the synthesized primers. The PCR product was cloned into a pCas9 plasmid removed the whole Cas9, trancRNA and spacer RNA sequences. The J23119-sgRNA sequence was amplified with PCR from pgRNA-bacteria (Addgene) and cloned into the newly prepared pCas9 vector. The new plasmid that can express Cas9 protein under the control of J23100 promoter and sgRNA under the control of J23119 promoter was named as pCas9-sgRNA. This plasmid also contained a chloromycetin gene under control of cat promoter. The various sgRNA sequences (annealed double-stranded oligonucleotides ended with BsaI sites) were then cloned into pCas9-sgRNA by using BsaI for simultaneously expressing Cas9 protein and the interested sgRNA in bacteria. The target sequences of sgRNA (plus PAM) used for the *in vivo* Cas9/sgRNA cutting were shown in Table 1.

**Table 1.**
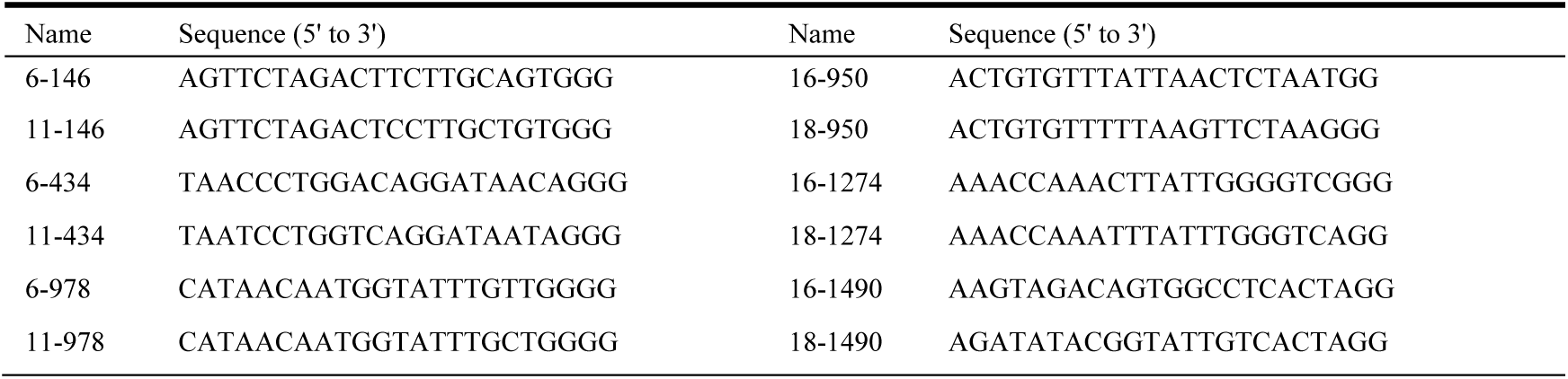
The target sequences of sgRNA (plus PAM) used for in vivo Cas9/sgRNA cutting

### *In vivo* cleavage of HPV16 and 18 L1 genes with Cas9/sgRNA

For the *in vivo* cutting of HPV L1 genes with Cas9/sgRNA, *E.coli* DH5α was firstly transfected with the plasmids cloned with HPV L1 gene and an ampicillin resistance gene (Amp^R^) under the control of Amp^R^ promoter. The transfected *E.coli* was selected on ampicillin agars and positive *E.coli* clones were confirmed by a clone PCR. The positive *E.coli* was then transfected with pCas9-sgRNA expressing various sgRNAs. The transfected *E.coli* was cultivated on agars with ampicillin plus chloromycetin overnight and imaged.

### Preparation of sgRNA

The sgRNA was synthesized by an *in vitro* transcription using T7 polymerase (New England Biolabs) according to the manufacturer’s instruction. In brief, sgRNA templates were generated by a three-round fusion PCR amplification using oligonucleotides listed in Table 2. The 1^st^-PCR was amplified with F1 and R (7 cycles); the 2^nd^-PCR was amplified with primers F2 and sgR using the 1^st^-PCR product as template (30 cycles), and the 3^rd^-PCR was amplified with primers F3 and sgR using the 2^nd^-PCR product as template (30 cycles). The purified 3^rd^-PCR products were used as template for preparing sgRNA by *in vitro* transcription. The in vitro transcription was performed by incubating the purified sgRNA template with T7 RNA polymerase (New England Biolabs) supplemented with rNTPs (New England Biolabs) overnight at 37 °C. The in vitro transcribed RNA was mixed with Trizol solution, and then successively extracted with chloroform and isopropanol, and precipitated with ethanol. Purified RNA was dissolved in the RNase-free ddH_2_O and quantified by spectrometry.

**Table 2.**
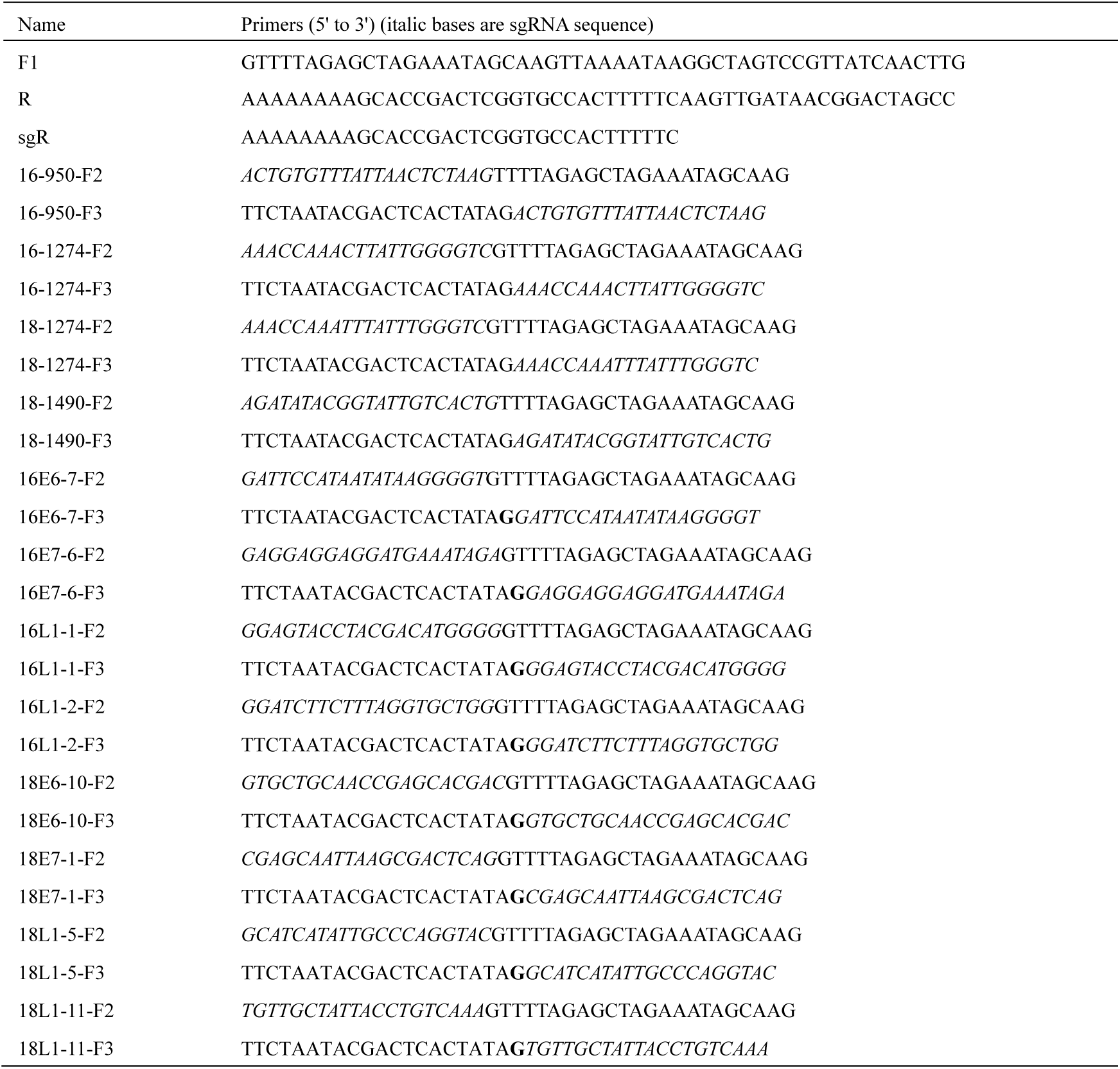
Oligonucleotides used to prepare the transcriptional template of sgRNA in this study

### Cleavage of HPV L1 genes cloned in plasmid with Cas9/sgRNA

Recombinant Cas9 protein was purchased from New England Biolabs (NEB). The Cas9 digestion reaction (30 μL) consisted of 1×Cas9 Nuclease Reaction Buffer, 1 μM Cas9 Nuclease (NEB), 300 nM sgRNA a (16-1274 or 18-1490; Table 2) and 300 nM sgRNA b (16-950 or 18-1274; Table 2) were firstly incubated at 25 °C for 10 min (this process was referred to as pre-assemble hereafter). 200 ng substrate DNA (L1 plasmid DNA linearized by AatII) mixed with above solution were incubated at 37 °C for 5 min. The reaction was mixed with 10×SDS-containing loading buffer (Takara) and run with 1.0% agarose gel.

### Detection of HPV 16 and 18 L1 genes cloned in plasmid with ctPCR

For preparing T adaptor, oligo oJW102 and oJW103 (Table 3) were dissolved in the Tris-HCl/EDTA/NaCl (TEN) buffer and mixed in the same molar. The mixture was heated at 95 °C for 5 min, and slowly cooled to room temperature. Plasmids cloned with the L1 genes of various HPV subtypes (200 ng) were cut by the Cas9 proteins in complex with a pair of sgRNAs targeting to HPV16 and HPV18 L1 genes. The plasmid (200 ng) was mixed with the pre-assembled Cas9/sgRNA complex that contained 1×Cas9 nuclease reaction buffer, 1 μM Cas9 nuclease, 300 nM sgRNA a (16-1274 or 18-1490; Table 2), and 300 nM sgRNA b (16-950 or 18-1274; Table 2) and incubated at 37 °C for 5 min. The digestion reaction (5 μL) was mixed with 5 μL premix Taq (Takara) and incubated at 72°C for 5 min. The A tailing reaction (10 μL) was mixed with 1×T4 Ligase Buffer, 830 nM T adaptor and 5 U T4 DNA Ligase and incubated at 22 °C for 5 min. The process of Cas9 cutting, A tailing and T adaptor ligation was concisely named as CAT hereafter. At last, the CAT-treated DNAs were amplified with tPCR by using a general primer (oJW102) annealing to T adaptor or a pair of general-specific primers (gs-primers) specific to the Cas9-digested HPV16 and HPV18 L1 genes. The tPCR reaction contained containing 10 μL SYBR Green (Bioer), 500 nM of a general primer (oJW102; Table 3) or 500 nM of each gs-primers specific to the L1 and E6-E7 genes of HPV16 and 18 (Table 3). The PCR program was as follows: 95°C 2 min, 30 cycles of 95°C 15 s, 60°C 30 s and 72°C 60 s, and 72°C 5 min. The reaction was run with 1.5% agarose gel.

**Table 3.**
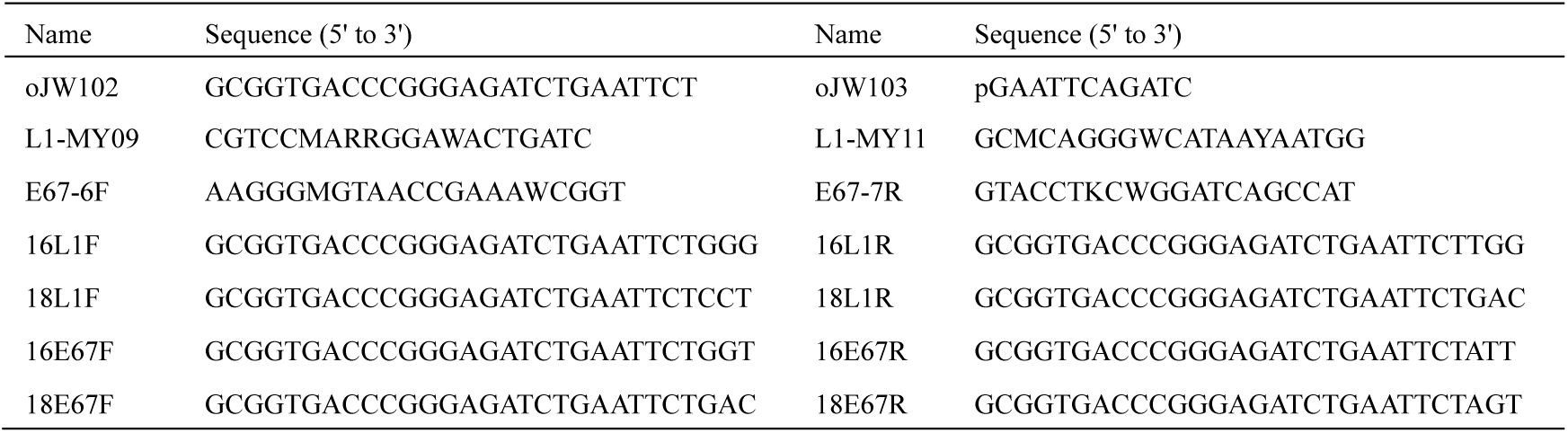
Oligonucleotides used as T adaptor and PCR primers in this study.

### Detection of HPV DNAs human cervical carcinoma cells with ctPCR

For the tPCR detection, PCR1 amplification of L1 or E6-E7 genes was carried out in a 20-μL tPCR reaction containing 10 μL premix primeSTAR Taq (Takara), 500 nM MY09 or E67-6F (Table 2), 500 nM MY11 or E67-7R (Table 2), various amounts of gDNA ( see figures) of three human cervical carcinoma cells (SiHa, HeLa and C-33a). The PCR program was as follows: 95°C 2 min, 35 cycles of 95°C 15 s, 60°C 30 s and 72°C 60 s, and 72°C 5 min. The PCR products were run with agarose gel. The PCR1 products (5 μL) were mixed with the pre-assembled Cas9/sgRNA complex that contained 1×Cas9 nuclease reaction buffer, 1 μM Cas9 nuclease, 300 nM sgRNA a (L1-1 or L1-5 for L1 gene; E6-7 or E6-10 for E6-E7 genes; Table 2), and 300 nM sgRNA b (L1-2 or L1-11 for L1 gene; E7-6 or E7-1 for E6-E7 genes; Table 2) and incubated at 37 °C for 5 min. The digestion reaction (5 μL) was mixed with 5 μL premix Taq (Takara) and incubated at 72°C for 5 min. The A tailing reaction (10 μL) was mixed with 1×T4 Ligase Buffer, 830 nM T adaptor and 5 U T4 DNA Ligase and incubated at 22 °C for 5 min. Finally the CAT-treated PCR1 products (1 μL) were amplified in a 20-μL tPCR reaction containing 10 μL SYBR Green (Bioer), 500 nM of each gs-primers specific to the L1 and E6-E7 genes of HPV16 and 18 (Table 3). The PCR program was as follows: 95°C 2 min, 30 cycles of 95°C 15 s, 60°C 30 s and 72°C 60 s, and 72°C 5 min. PCR program was ere run on a tPCR machine, 9700 (ABI). The PCR2 products were run with 1.5% agarose gel.

For the qPCR detection, qPCR1 amplification of L1 or E6-E7 genes was carried out in a 20-μL qPCR reaction containing 10 μL 2×Sybr Green Master Mix (Yeasen), 500 nM MY09 or E67-6F (Table 2), 500 nM MY11 or E67-7R (Table 2), various amounts of gDNA of three human cervical carcinoma cells (SiHa, HeLa and C-33a). The qPCR program was as follows: 95°C 10 min, 40 cycles of 95°C 15 s, 60°C 30 s and 72°C 1 min. The qPCR1 products (2 μL) were mixed with the pre-assembled Cas9/sgRNA complex that contained 1×Cas9 nuclease reaction buffer, 1 μM Cas9 nuclease, 300 nM sgRNA a (L1-1 or L1-5 for L1 gene; E6-7 or E6-10 for E6-E7 genes; Table 2), and 300 nM sgRNA b (L1-2 or L1-11 for L1 gene; E7-6 or E7-1 for E6-E7 genes; Table 2). The reaction was incubated at 37 °C for 5 min. The digestion reaction (5 μL) was mixed with 5 μL premix Taq (Takara) and incubated at 72°C for 5 min for A tailing. The A tailing reaction (10 μL) was mixed with 1×T4 Ligase Buffer, 830 nM T adaptor and 5 U T4 DNA Ligase and incubated at 22 °C for 5 min. Finally the CAT-treated qPCR1 products (1 μL) were amplified in a 20-μL qPCR reaction containing 10 μL 2×Sybr Green Master Mix (Yeasen), 500 nM of each gs-primers specific to the L1 and E6-E7 genes of HPV16 and 18 (Table 3). The PCR program was as follows: 95°C 10 min, 40 cycles of 95°C 15 s, 60°C 30 s and 72°C 1 min. PCR program was run on a real-time PCR machine, StepOne Plus (ABI). The qPCR1 and qPCR2 products were also sometimes run with 1.5% agarose gel for further checking specificity in addition to melting curve.

## Results

### Cleavage of HPV 16 and 18 L1 genes with Cas9/sgRNA

To preliminarily explore if the Cas9/sgRNA system could be used to specifically discriminate subtypes of HPVs, we firstly performed an *in vivo* cutting assay. In this assay, *E.coli* DH5α was firstly transfected with a HPV L1 plasmid that has ampicillin resistance. The positive cells were then transfected with a Cas9/sgRNA-expressing plasmid that has Chloromycetin resistance. After overnight cultivation, agar plates were imaged. We found that the *E.coli* with a HPV L1 plasmid was always specifically killed by Cas9 nuclease guided by sgRNA specific to the HPV L1 gene (Fig.1). These results revealed that the designed sgRNA could specifically target to its target and the Cas9/sgRNA could be used to type HPV subtypes.

**Fig.1.**
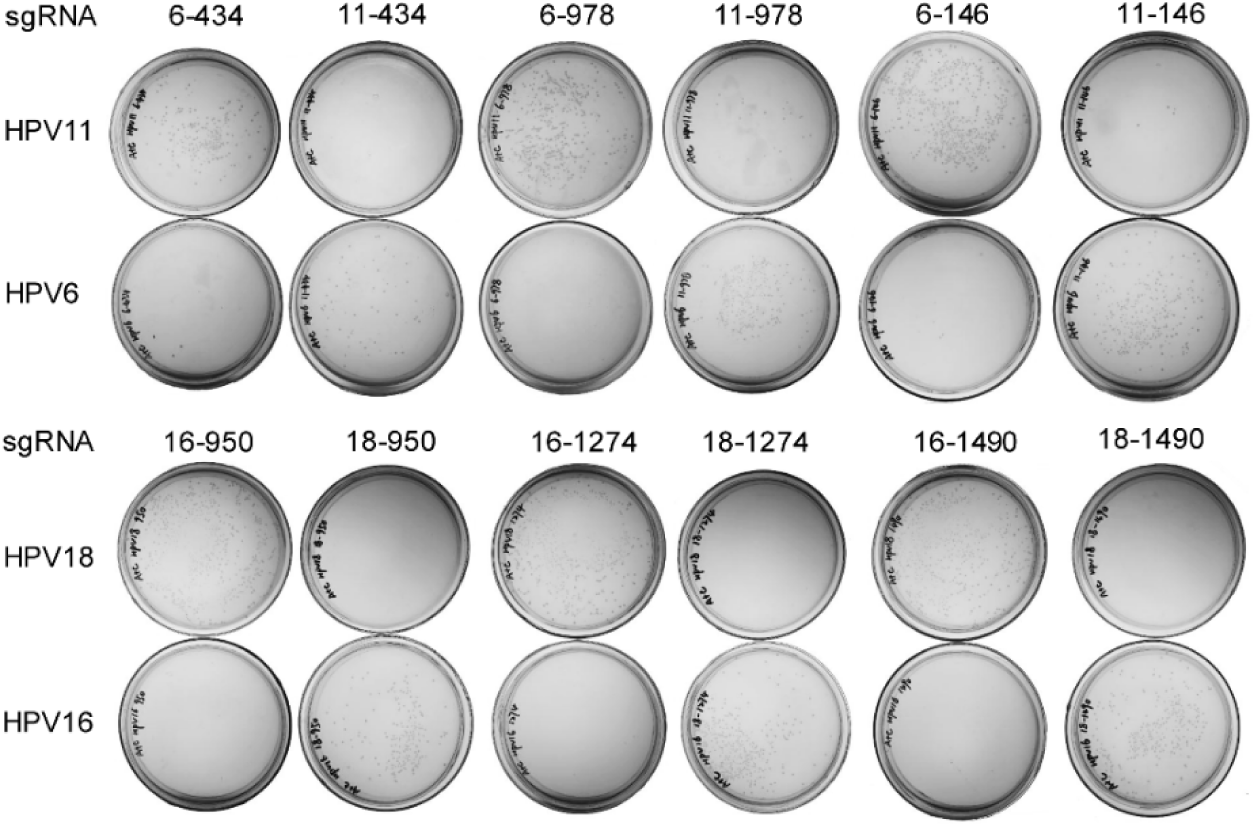
*In vivo* cutting of HPV L1 genes with Cas9/sgRNA. The *E.coli* DH5α was firstly transfected with HPV L1 plasmid and the selected Amp-resistant cells were then transfected with Cas9/sgRNA plasmid. The transfected cells were cultivated on agars with ampicillin plus chloromycetin overnight and imaged.

### Cleavage of HPV L1 genes cloned in plasmid with Cas9/sgRNA

With the high-efficiency *in vivo* cutting of HPV L1 genes by Cas9/sgRNA, we conceived if the kind of specific DNA cutting by the sgRNA-guided Cas9 could be used to detect and type DNA *in vitro*. We thus performed the *in vitro* cutting assays. We firstly linearized the plasmids cloned with the HPV16 and HPV18 L1 genes with a restriction endonuclease, AatII, which produced linear DNA fragments ended with a 4-base 3′ overhang. We cut the linearized HPV16 and HPV18 L1 plasmid DNAs with Cas9 nuclease in complex with the sgRNAs specific to HPV16 and HPV18 L1 genes (Table 2). The results indicated that the HPV16 and HPV18 L1 genes could be specifically targeted by their corresponding sgRNA and cut by the guided Cas9 nuclease (Fig.2). This means that the *in vitro* specific DNA cutting by Cas9/sgRNA could be used to detect and type DNA.

**Fig.2.**
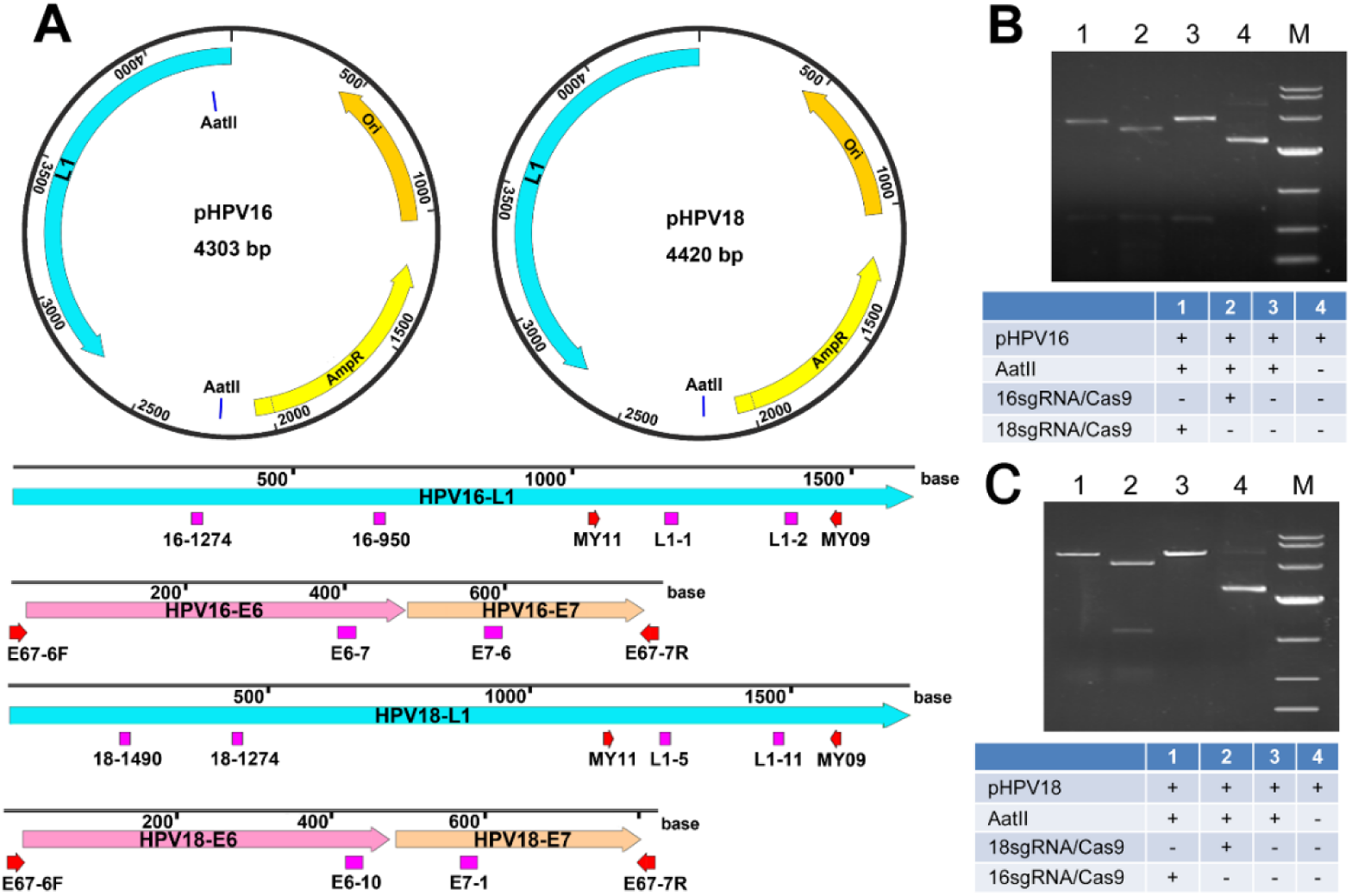
Cas9/sgRNA cleavage of HPV16 and HPV18 L1 genes. (A) HPV L1 plasmids and locations of sgRNA targets and universal PCR primers in the L1 and E6-E7 genes of HPV16 and HPV18. (B) Cas9/sgRNA cleavage of HPV16 L1 gene using sgRNA 16-1274 and 16-950. (C) Cas9/sgRNA cleavage of HPV18 L1 gene using sgRNA 18-1490 and 18-1274. The Cas9 protein was in complex with the sgRNAs specific to HPV16 or 18 L1 genes and used to cut the linearized HPV16 or HPV18 L1 plasmids (A). The DNAs were run with agarose gel.

### Detection of HPV 16 and 18 L1 genes with ctPCR

To detect and type DNA with Cas9/sgRNA, we designed a CRISPR-typing PCR (ctPCR) method for realizing this end. In this method, a target DNA was firstly cut with a pair of sgRNAs specific to the interested target DNA. The Cas9/sgRNA-cut DNA was then tailed with an adenine (A) and ligated with a T adaptor. We concisely named the process of Cas9 cutting, A tailing and T adaptor ligation as CAT hereafter. Finally, the CAT-treated DNA was amplified with PCR by using a constant primer annealing to T adaptor (Fig.3A). We used the method detected HPV16 and HPV18. The results revealed that the expected HPV16 target DNA was successfully and specifically detected by the method (Fig.3B), while there was an additional nonspecific DNA fragment when detecting HPV18 with the method (Fig.3C). In order to improve the detection specificity, we added three specific nucleotides at the 3′ end of the constant primer annealing to T adaptor. We named this kind of primer as general-specific primer (gs-primer). We prepared a pair of gs-primers for HPV16 and HPV18 according to the cutting sites of HPV16 and HPV18 sgRNAs. We then amplified the CAT-treated DNA with a pair of gs-primers. As a result, we found that the target HPV16 and HPV18 DNAs were specifically detected by the improved ctPCR method (Fig.3, B and C).

**Fig.3.**
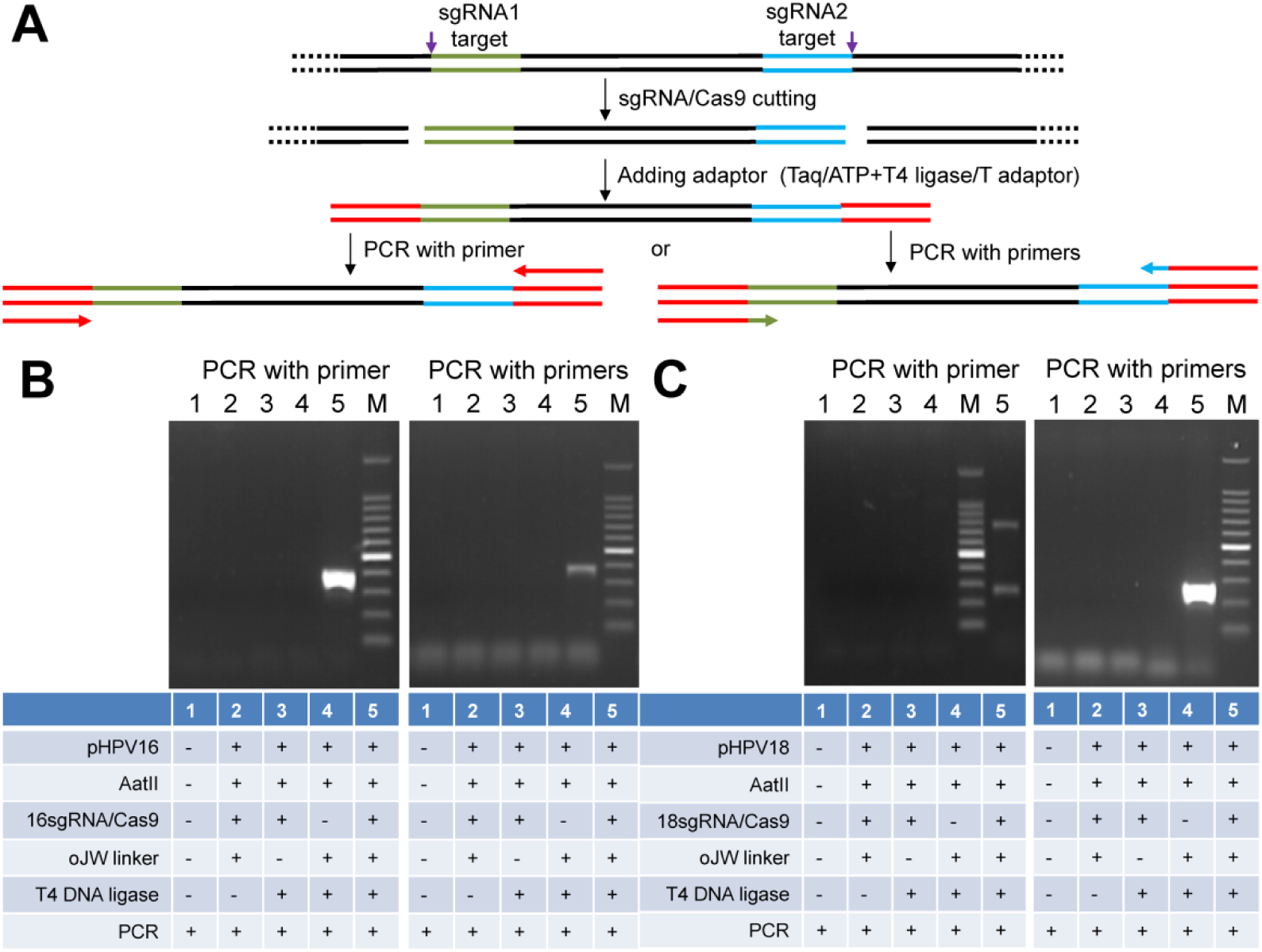
Detection of HPV L1 gene with ctPCR by using different primers. (A) A schematic of the experimental procedures for detecting and typing HPV DNA with ctPCR by using different primers. (B) Detection of HPV16 L1 gene by ctPCR with different primers. (C) Detection of HPV18 L1 gene by ctPCR with different primers. The final PCR products were run with agarose gel. Primer, a general primer complementary to the constant T adaptor (oJW linker) used in ctPCR detection; Primers, a pair of primers complementary to constant T adaptor and 3 nucleotides at the end of Cas9-cut product (named as general-specific primers, gs-primers).

### Sensitivity of ctPCR detection of HPV16 and HPV18 L1 gene

We next investigated the detection sensitivity of L1 gene with ctPCR. Different amount of HPV16 and HPV18 L1 gene plasmids was respectively cut with the Cas9 nuclease in complex with pair of sgRNAs targeting to the HPV16 and HPV18 L1 genes. The cut DNA was tailed with A and ligated with T adaptor. Then the CAT-treated DNA was amplified by PCR2 with the corresponding pairs of gs-primers and the PCR products were detected with agarose gel electrophoresis. The results revealed that ctPCR showed high amplification efficiency and sensitivity. It was found that as little as 5 ng of CAT-treated HPV18 L1 plasmid DNA was detected by tPCR-based ctPCR (Fig.4). In addition, when the CAT-treated HPV18 L1 plasmid DNA was amplified by qPCR, as little as 10000-fold diluted cat-treated HPV18 L1 plasmid DNA could be detected. These data suggested that the CAT treatment had high efficiency.

**Fig.4.**
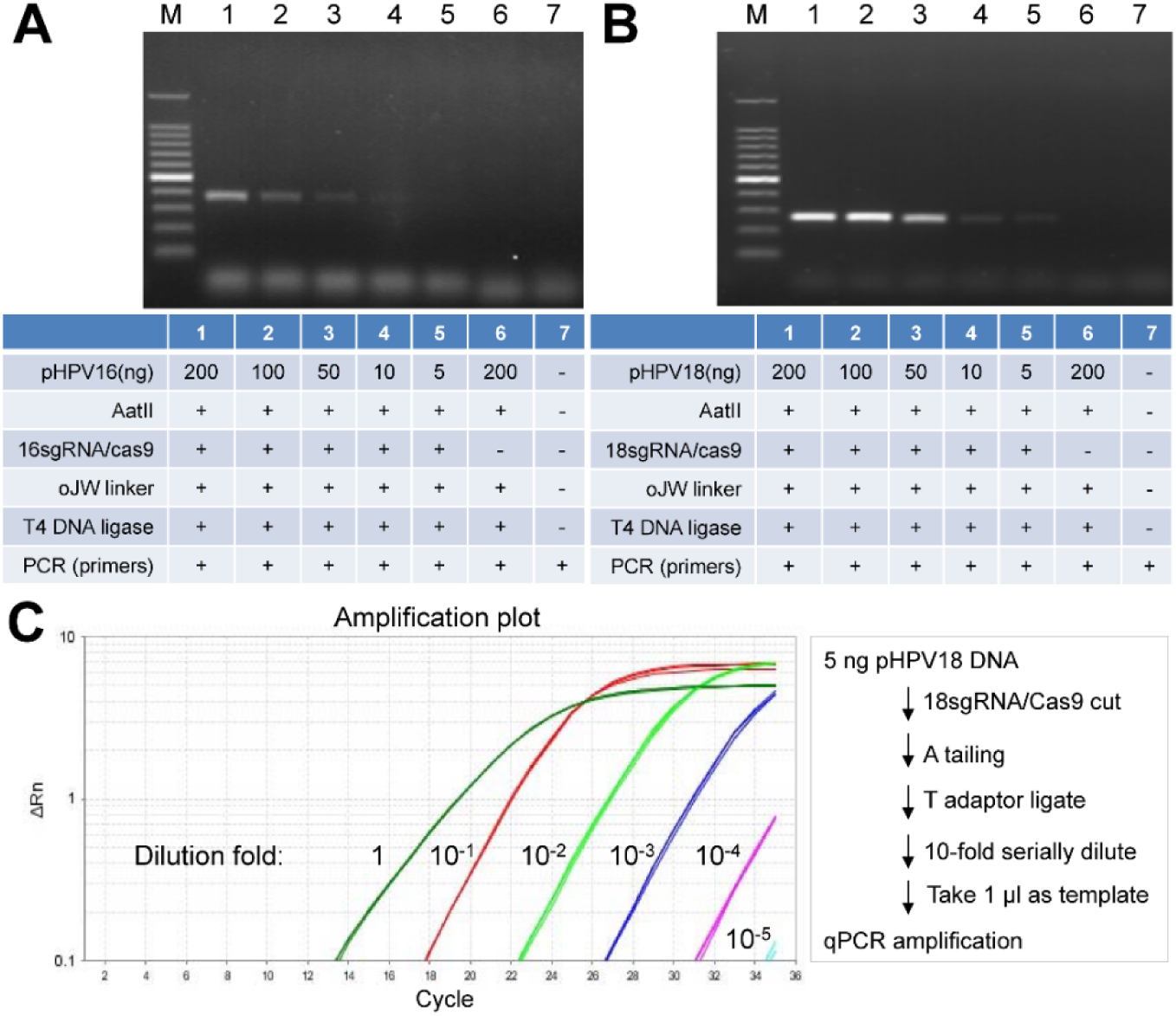
Sensitivity of ctPCR detection of HPV 16 or 18 L1 genes. (A) Detection of HPV 16 L1 gene with tPCR-based ctPCR. (B) Detection of HPV 18 L1 gene with tPCR-based ctPCR. (C) Detection of HPV 18 L1 gene with qPCR-based ctPCR.**Fig.8** Detection of HPV18 L1 gene in HeLa cells with qPCR-based ctPCR. (A) Detecting the HPV18 L1 gene in the HeLa gDNA with qPCR1 by using universal primers MY09/11. (B) Typing the HPV18 L1 gene with ctPCR. The final ctPCR products were also run with agarose gel.

### Detection of HPV 16 or 18 L1 genes in 13 HPV subtypes with ctPCR

In order to further verify the specificity of ctPCR, we digested the L1 genes of 12 HPV subtypes with the Cas9 protein in complex with the sgRNAs of HPV16 or HPV18. The digested DNAs were then tailed with A and ligated with T adaptor. The CAT-treated DNAs were amplified with tPCR by using gs-primers. Finally, the tPCR products were run with agarose gel. The results revealed that the L1 genes of HPV16 and HPV18 were specifically detected by ctPCR from other 11 HPV subtypes (Fig.5). Two highest risk HPV subtypes, HPV16 and HPV18, were discriminated from other 10 high-risk HPV subtypes.

**Fig.5.**
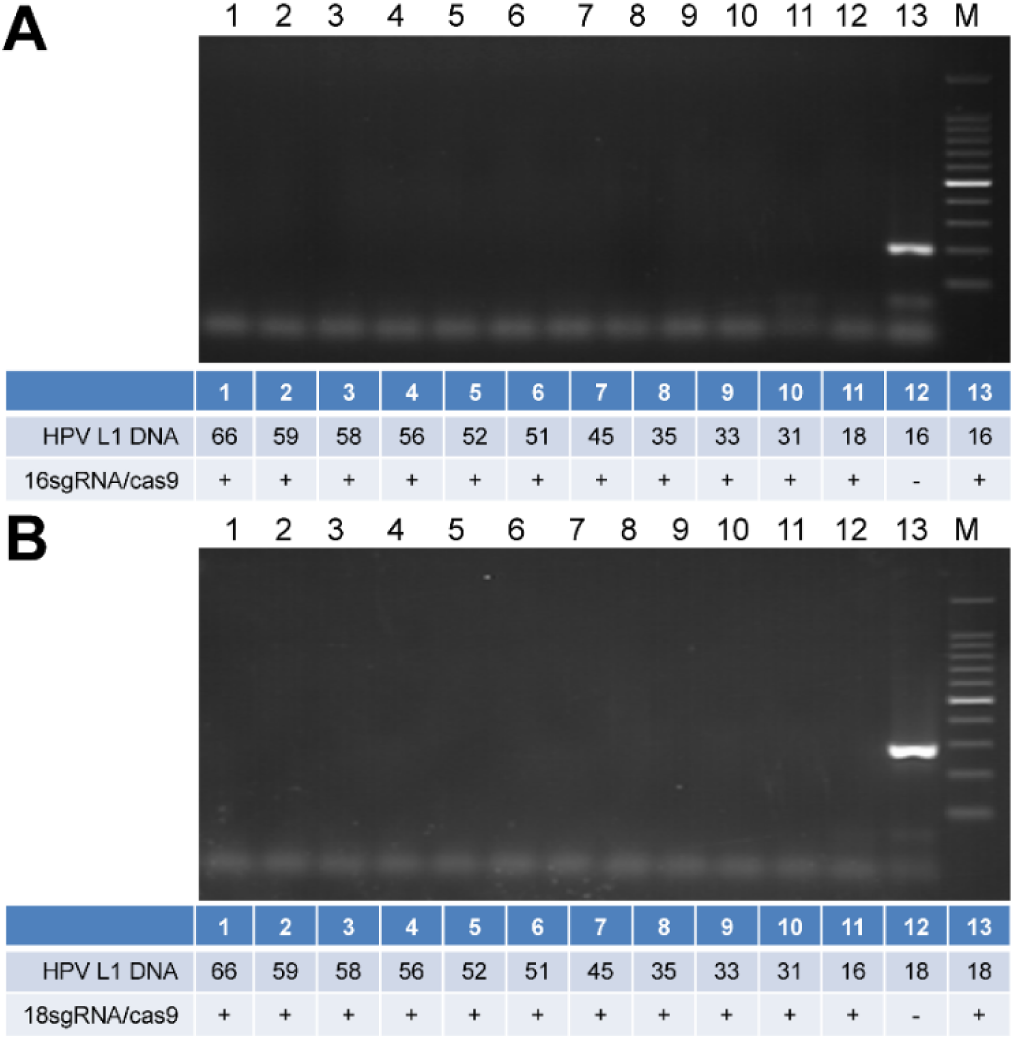
Detection of HPV 16 or 18 L1 genes in thirteen HPV subtypes with ctPCR. (A) Detection of HPV 16 L1 gene in thirteen HPV subtypes. (B) Detection of HPV 18 L1 gene in thirteen HPV subtypes. The final ctPCR products were run with agarose gel.

### Detection of HPV genes in the cervical carcinoma cells with ctPCR

Despite the L1 genes could be detected by ctPCR, the L1 genes together with its host plasmid is a relative simple DNA sample to ctPCR detection. However, the HPV clinical detection uses the complex genomic DNAs (gDNAs) of cells. To check if the ctPCR technique could be used to detect HPV in gDNAs, we next tried to detect the L1 and E6-E7 genes of HPVs in the gDNAs of human cervical carcinoma cells according to a two-round PCR strategy (Fig.6A). To this end, we firstly prepared the gDNAs from three different human cervical carcinoma cell lines, HeLa, SiHa and C-33a. We then amplified the L1 gene by using a pair of universal primers, MY09 and MY11, which was previously designed to amplified the L1 genes of various HPV subtypes. As a result, the L1 gene was successfully amplified from the HeLa and SiHa gDNAs but not from the C-33a gDNA (Fig.6, B and C). Because no such universal primers were reported for amplifying the E6 and E7 genes, we newly designed a pair of such universal primers, E67-6F and E67-7R, for amplifying the E6-E7 genes of various HPV subtypes. We then amplified the E6-E7 genes by using the pair of primers. The results revealed that the E6-E7 genes could be amplified from the HeLa and SiHa gDNAs but not from the C-33a gDNA (Fig.6, B and C). We called this first-round PCR as PCR1 (Fig.6A). We next digested the L1 and E6-E7 qPCR1 products with the Cas9 nuclease in complex with pairs of sgRNAs specific to the L1 and E6-E7 genes of HPV16 and HPV18. After the Cas9-cut PCR1 products were tailed with A and ligated with T adaptor, the CAT-treated DNA was amplified with pairs of gs-primers specific to the L1 and E6-E7 genes of HPV16 and HPV18. We called this second-round PCR as PCR2 (Fig.6A). Although no L1 and E6-E7 genes were detected from the C-33a gDNA by PCR1, we still treated the PCR1 products of the C-33a gDNA with CAT and amplified with PCR2. However, no L1 and E6-E7 genes of HPV16 and 18 were detected (Fig.6, B and C). These results are in agreement with the previous reports that HeLa and SiHa are HPV18- and HPV16-positive cells^24,25^, respectively, and C-33a is a HPV-negative cell ^26^.

**Fig.6.**
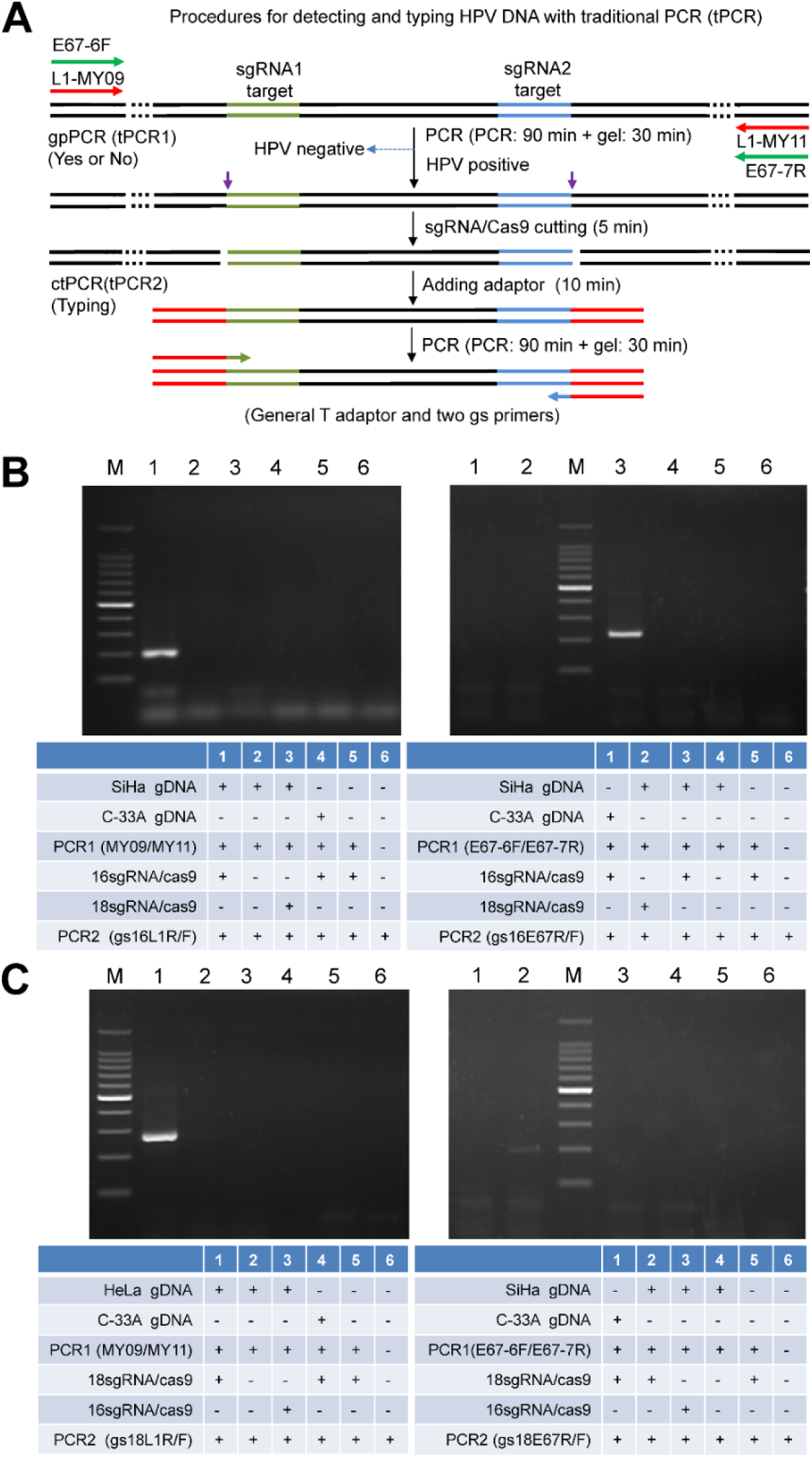
Detection of HPV16 and HPV 18 genes in cervical carcinoma cells with tPCR-based ctPCR. (A) Schematic show of procedures for detecting and typing HPV DNA with tPCR-based ctPCR. (B) Detecting the HPV16 L1 and E6-E7 genes in the SiHa gDNA (200 ng) with ctPCR. (C) Detecting the HPV18 L1 and E6-E7 genes in the HeLa gDNA (200 ng) with ctPCR. The C-33a gDNA (200 ng) was detected as a negative control and a mimic detection of no DNA was used as a blank control. The final ctPCR products were run with agarose gel.

Despite the L1 and E6-E7 genes could be detected with ctPCR from the gDNAs of human cervical carcinoma cells, such tPCR-based ctPCR detection is unfavorable to its clinical application due to cumbersome gel electrophoresis. We therefore investigated if the ctPCR detection could be realized with a similar two-round qPCR procedure (Fig.7A). As expected, both L1 and E6-E7 genes were easily amplified by qPCR1 from gDNAs of HeLa and SiHa cells (Fig.7, B and C). We next treated the qPCR1 products with CAT and amplified with qPCR2 by using gs-primers. As expected, the HPV16 and HPV18 L1 and E6-E7 genes were successfully detected in the gDNA of SiHa and HeLa cells, respectively (Fig.7, B and C). This means that the gradients of qPCR1 did not interfere the followed CAT treatments and the gradients of CAT treatments also did not interfere the followed qPCR2. This greatly simplified the qPCR-based ctPCR detection due to no DNA purification step needed.

**Fig.7.**
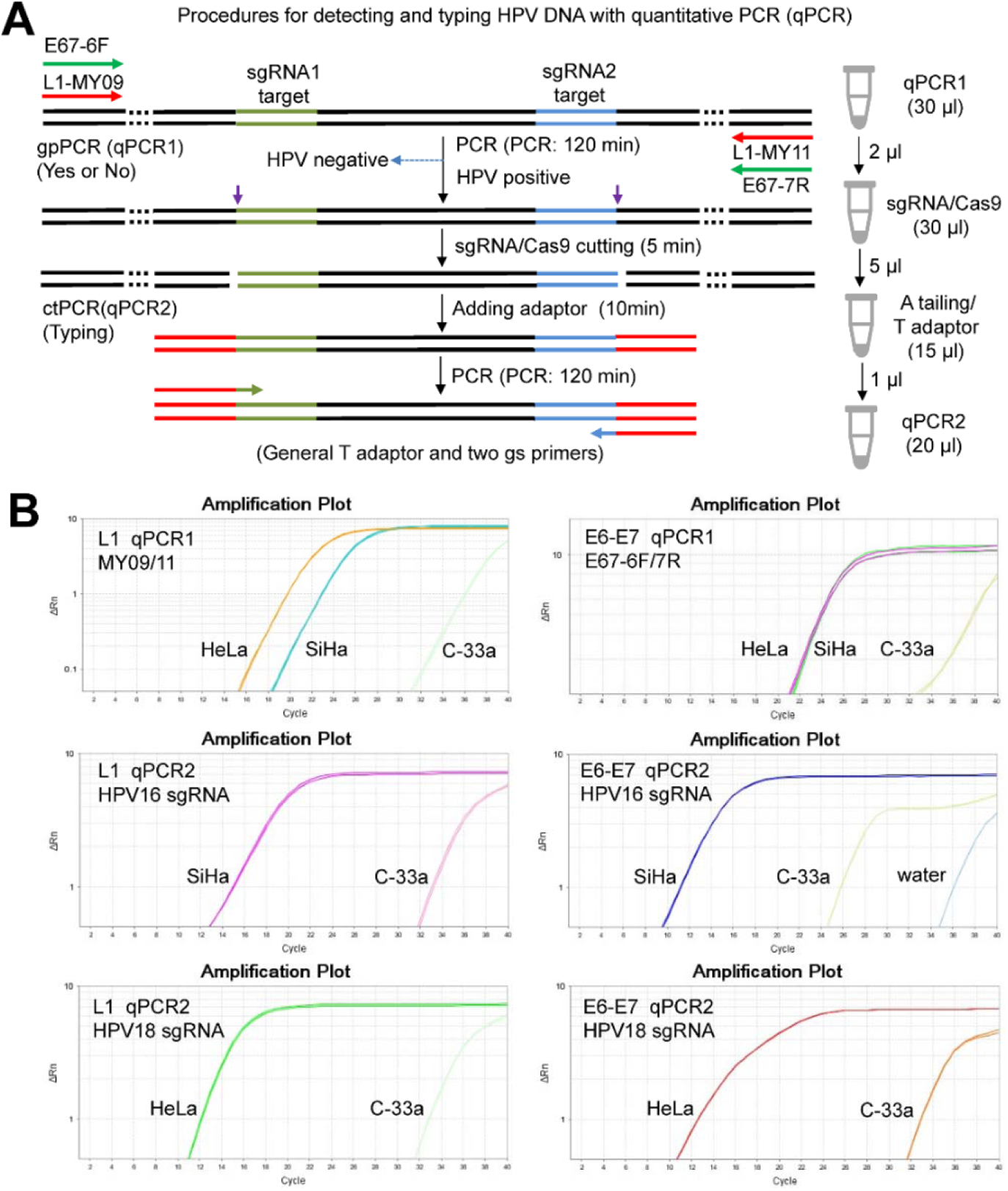
Detection of HPV16 and HPV 18 genes in cervical carcinoma cells with qPCR-based ctPCR. (A) Schematic show of procedures for detecting and typing HPV DNA with qPCR-based ctPCR. The reaction volumes of each step and the solutions used to the next step was shown (right). (B) Detecting the HPV16 L1 and E6-E7 genes in the three human cervical carcinoma cell lines, HeLa, SiHa and C-33a. Q-PCR1 used 200 ng gDNA of each cell line as template.

In order to check the detection specificity of ctPCR, we used as many as 200 ng gDNA as template of PCR1 to detected HPV L1 and E6-E7 genes in the tPCR- and qPCR-based ctPCR detection. We next checked the detection sensitivity of both qPCR and qPCR2. To this end, we amplified the HPV18 L1 gene from various amount of HeLa gDNAs with qPCR1. The results indicated that the HPV18 L1 gene could be amplified qPCR1 from various amount of HeLa gDNAs (Fig.8A). Especially, the HPV18 L1 gene was amplified by qPCR1 from as little as 0.005 ng gDNA (Fig.8A). In addition, even the CAT-treated qPCR1 product of 0.005 ng gDNA template was diluted 1000 fold (10^-3^), qPCR2 could amplify HPV18 L1 gene using 1 μL of the diluted solution (Fig.8B). These results indicated that both PCR1 and PCR2 can be realized with qPCR in high sensitivity, suggesting that the qPCR-based ctPCR is favorable to clinical application.

**Fig.8.**
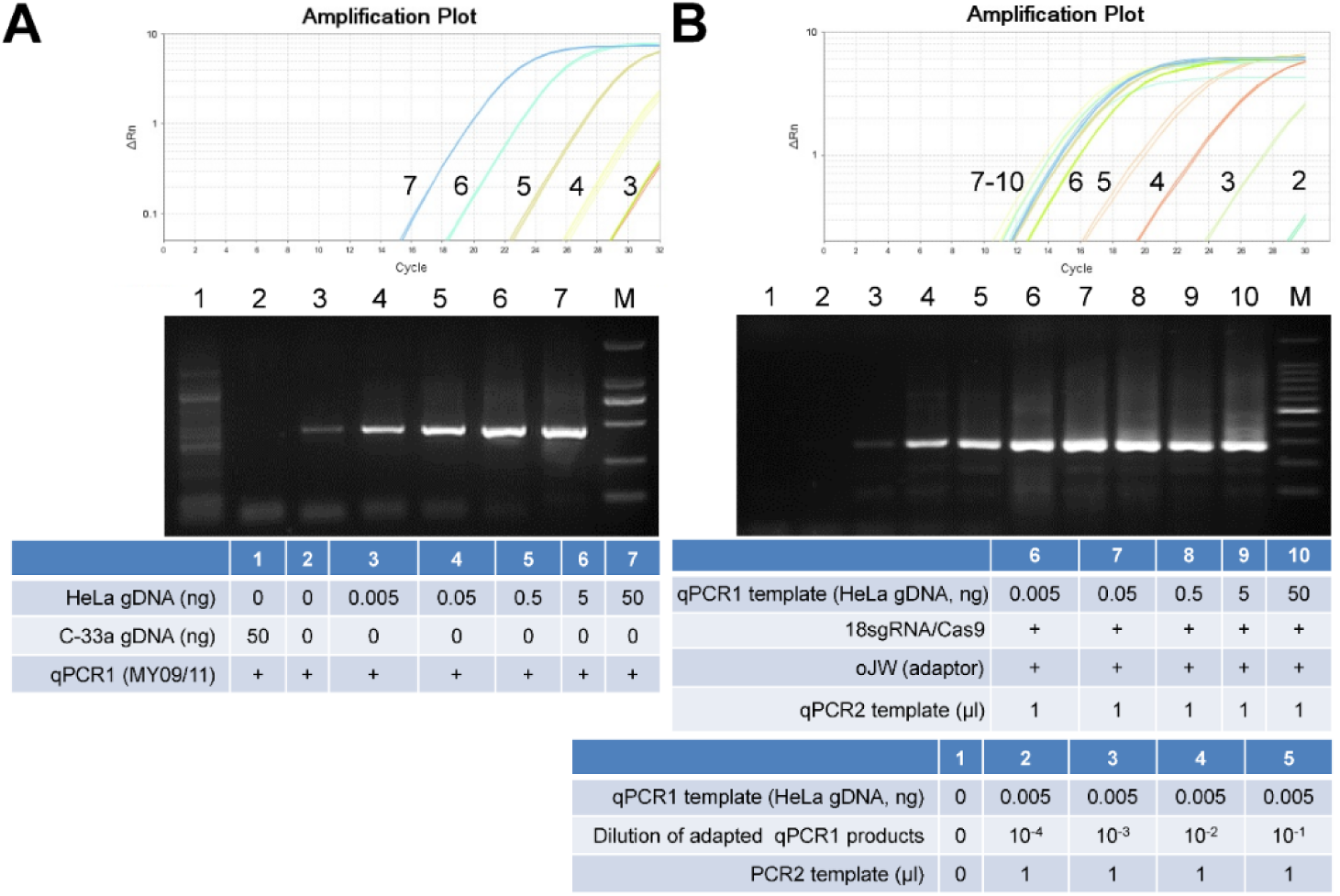
Detection of HPV18 L1 gene in HeLa cells with qPCR-based ctPCR. (A) Detecting the HPV18 L1 gene in the HeLa gDNA with qPCR1 by using universal primers MY09/11. (B) Typing the HPV18 L1 gene with ctPCR. The final ctPCR products were also run with agarose gel.

## Discussion

HPV is a kind of dsDNA virus that is the pathogen of cervical, anogenital, and other cancers ^27-29^. There are around 100 HPV subtypes with different genetic variations ^30^. According to the oncogenic potential, HPVs are divided into high- and low-risk HPVs ^31^. The most common world-wide high-risk HPVs are HPV16 and HPV18 ^32-34^, which lead to approximately 70% of cervical cancers ^35^. Other high-risk HPVs include HPV31, 33, 35, 39, 45, 51, 52, 56, 58, 59, 66, and 68 ^36^. The low-risk HPVs include HPV6, 11, 40, 42, 43, 44, 61, and 81 ^37,38^. Due to its carcinogenesis, HPV detection is at present widely performed in cervical cancer diagnosis and routine women health examination by mainly using various PCR-based techniques ^34,35^. For example, the Roche cobas HPV testing (cobas 4800 or cobas 6800/8800 Systems) is clinically validated for HPV primary screening. The cobas HPV assays provide specific genotyping information for HPV16 and HPV18, while simultaneously reporting the 12 other high-risk HPV types as a pooled result, all in one test and from one patient sample. Because of highly variable genotypes, HPV is an ideal material for testifying the nucleic acid detection and genotyping methods.

The L1 gene has been widely used to detect and type HPVs. In this study, we firstly designed sgRNAs for detecting HPV L1 genes because we have a set of plasmids cloned with L1 genes of various HPV subtypes (Fig.5). We performed the *in vivo* and *in vitro* cutting assays of HPV L1 genes with Cas9/sgRNA. We also verified the ctPCR method by detecting the HPV16 and HPV 18 L1 genes, which helped us to improve the ctPCR method by introducing gs-primers. We also finally detected two high-risk HPVs, HPV16 and HPV18, in two cervical carcinoma cells, HeLa and SiHa, by detecting the L1 genes with ctPCR. However, it has been reported that HPV can lose its L1 gene when integrating into host cell genome, which can thus lead to false-negative detection ^29,39,40^. Additionally, it was found that HPV18 integrated into the human genome in almost all HPV18-positive cervical cancers, and HPV16 integrated into the human genome in about 60% of HPV16-positive cervical cancers ^29^. Therefore, HPV detection is increasingly relying on the carcinogenesis gene E6/E7 ^41^, which can avoid missed detection because E6/E7 always exists after integration. Therefore, the E6/E7 genes can be used as the reliable targets of HPV detection. This study thus also detected E6/E7 genes when detecting HPV16 and HPV18 in three human cervical cancer cell lines, HeLa, SiHa and C-33a. The results demonstrated that the two highest-risk HPVs, HPV16 and HPV18, could be detected from SiHa and HeLa cells, respectively; however, both HPVs were not detected in C-33a. This is in agreement with the facts that HeLa is a HPV18-positive cell ^24^, SiHa is a HPV16-positive cell ^25^, but C-33a is a HPV-negative cell ^26^.

The partial L1 missing in the integrated HPV DNA was also observed by this study. In this study, we initially designed a pair of sgRNAs for both HPV16 and HPV18 L1 genes (16-1274/16-950; 18-1490/18-1274; Fig.2). We performed the *in vivo* and *in vitro* cutting assays of L1 plasmid using these sgRNAs (Fig.2 to 5). However, when detected HPV18 in HeLa cells using sgRNA 18-1490/18-1274, we found that no DNA could be amplified by ctPCR. Because it was reported that the L1 gene can be detected in the HeLa cells by using universal primers, MY09/11 ^42^, we thus designed a new pair of sgRNAs for HPV16 and HPV18 L1 genes (L1-1 and L1-2 for HPV16; L1-5 and L1-11 for HPV18; Table 2; Fig.2), which locate in the L1 region that can be amplified by primer MY09/11. We re-amplified L1 gene in three cervical carcinoma cell lines. As a result, the HPV L1 gene was found in the HeLa and SiHa. We then typed HPV L1 gene of two cells with ctPCR, which demonstrated that HPV16 and 18 L1 genes were present in SiHa and HeLa cells, respectively. These data suggest that the L1 region that was targeted by the initially designed sgRNAs was missed in the integrated HPV DNA. Nevertheless, the *in vivo* cutting and *in vitro* ctPCR detection of HPVs with two pairs of sgRNAs suggest that the multiple subtype-specific sgRNA can be easily designed due to the wide presence of PAMs in genome and high targeting specificity of Cas9/sgRNA system. It means that ctPCR relying on sgRNAs has higher genotyping capability than the traditional PCR that depends on the specific primers.

In this study, when detecting HPV16 and HPV18 DNAs in cervical carcinoma cell lines, we used a two-round PCR strategy. The first round PCR amplified L1 gene with universal primer MY09/11 or E6-E7 genes with universal primer E67-6F/7R. The PCR products were cleaved with Cas9/sgRNA, tailed with adenine (A), and ligated with a constant T adaptor. Then the second round of PCR was performed with a general primer or a pair of general-specific primers (gs-primers). Therefore, the first round PCR was used to amplify the HPV DNA for judging if the sample was infected by HPVs. The second round PCR, together with Cas9/sgRNA cutting, A tailing and T adaptor ligation, was named as CRISPR-typing PCR (ctPCR), which was used to type or discriminate the HPV subtypes infected the sample. Due to the high sensitivity of PCR amplification, the detection limit was warranted by the first round PCR (PCR1). Additionally, PCR1 also provided enough target DNA for the followed ctPCR.

DNA typing is critical for the specific DNA detection, which is especially useful for the discriminatively detection of virus subtypes and polymorphic DNAs. In this study, to guarantee the specificity of ctPCR detection, we adopted two strategies. One is to design two highly specific sgRNA commonly targeting to the detected DNA type. The other is to use a pair of gs-primers in ctPCR. Although the off-target of Cas9/sgRNA system is now limiting its application in human gene therapy, this disadvantage did not affect the ctPCR detection due to the double sgRNA cutting in a short DNA region. Additionally, the distant off-targets beyond the PCR amplification limit also did not affect the ctPCR. Even the off-targets cutting produced a fragment that is in the range of PCR amplification, these off-targets can be prevented from ctPCR amplification by the gs primers. The gs-primers further guarantee the detection specificity of ctPCR on the base of a pair of specific sgRNAs. Therefore, this study demonstrated that ctPCR could specifically detect the L1 and E6-E7 genes of the two highest risk HPVs (HPV16 and HPV18) integrated in the complicated genomic DNAs of two human cervical carcinoma cell lines, HeLa and SiHa.

Real-time PCR has already become a widely popularized DNA detection tool in clinics. For the clinical application purpose, this study investigated the feasibility to realize ctPCR detection with qPCR. It was found that the Cas9/sgRNA cutting could be directly performed with the qPCR1 solution, indicating that the gradients of qPCR1 reaction did not interfere the next Cas9/sgRNA cutting. Similarly, the gradients of Cas9/sgRNA reaction did not interfere the subsequent A tailing and T adaptor ligation, which also did not interfere the final qPCR2. Therefore, the whole ctPCR detection process is free of DNA purification and just realized by simple three-steps solution transfer (Fig.7A). This kind of compatibility among various biochemical reactions of different functions make the ctPCR detection easily and rapidly realizable in clinical detection. In addition, the qPCR-based ctPCR also increased the detection limit.

Clearly, the test cycle of ctPCR detection is mainly dependent on two round of PCR. In this study we have in fact optimized the CAT steps. We thus provided an optimized CAT treatment program of shortest time (Fig.6 and Fig.7). We found that five minutes was enough for Cas9/sgRNA to cut its targets in the ctPCR detection, indicating the high *in vitro* cutting efficiency of Cas9/sgRNA ^43^. We also found that five minutes was enough for A tailing and T adaptor ligation, respectively. Therefore, the whole CAT treatment process can be finished as few as 15 min. In fact, PCR1 contributes to the high efficiency of Cas9/sgRNA cutting due to enriching the detected targets. In this study, we have ever tried detecting HPV18 in the HeLa gDNA as we detected L1 plasmid; however, it failed, even after cutting large amount of gDNA (1 μg) for a long time (2 h). This agrees with a recent finding that Cas9/sgRNA complex spends as long as 6 hours to find its target in the *E.coli* genome (about 4 million base pairs) ^44^ Therefore, it will take Cas9/sgRNA complex very long time to find low-copy targets in a highly complicated DNA context such as human gDNA. The case can be further deteriorated by the fact that Cas9 is a single-turnover enzyme in DNA cutting ^43^. The problem may be addressed by using more Cas9/sgRNA complex and long-time digestion. However, a time-consuming and costly detection is prohibitive to clinical application. Nevertheless, a PCR1-free ctPCR should be effective to high-copy targets as we detected L1 plasmid.

It should be noted that this study focused on validating the ctPCR method in proof-of-concept by using HPV as a convenient available experimental material. The method could also be used to detect other interested DNA. This study employs HPV DNA as DNA target for ctPCR detection. The results indicate that ctPCR can detect and type HPV DNAs. It was found that ctPCR could detect the HPV L1 gene in as little as 0.005 ng gDNA from cervical carcinoma cell.

## Conclusion

This study developed a new method for detecting target DNA based on Cas9 nuclease, which was named as ctPCR, representing the Cas9/sgRNA-typing PCR. This method was verified by successfully detecting the L1 genes of two high-risk HPVs (HPV16 and HPV18) from 13 HPV subtypes. This method was also verified by successfully detecting the L1 and E6/E7 genes of two high-risk HPVs (HPV16 and HPV18) in three cervical carcinoma cells (HeLa, SiHa and C-33a). This study demonstrated that ctPCR has high specificity and sensitivity. This study also demonstrated that ctPCR detection could be realized by a simple two-round qPCR, making ctPCR applicable to clinical diagnosis. By using the widely available qPCR machines, the whole ctPCR detection process can be finished in as few as 3 to 4 hs. Therefore, ctPCR should be useful in DNA detection and genotyping in the future.

## Acknowledgments

This work was supported by the grants from the National Natural Science Foundation of China (Grant 61571119).

## Author contributions

J.K.W. conceived the idea and designed and instructed the study. Q.W., B.B.Z., and X.H.X, performed experiments. F.F.L. prepared cells. J.K.W. wrote the paper.

## Data availability statement

Not applicable.

